# Multi-Omics Analysis Of Antiviral Interactions Of *Elizabethkingia anophelis* And Zika Virus

**DOI:** 10.1101/2023.12.27.573398

**Authors:** S Omme, J Wang, M Sifuna, J Rodriguez, NR Owusu, M Goli, P Jiang, P Waziha, J Nwaiwu, CL Brelsfoard, A Vigneron, AT Ciota, LD Kramer, Y Mechref, MG Onyango

**Author notes:** Corresponding author: MGO. **Email addresses:** OS, WJ, SM, RJ, ONR, GM, JP, WP, NJ, CLB, VA, CAT, KLD, MY.

## Abstract

**Background:** The microbial communities residing in the mosquito midgut play a key role in determining the outcome of mosquito pathogen infection. *Elizabethkingia anophelis*, originally isolated from the midgut of *Anopheles gambiae*, has drawn much attention due to its close association with *Aedes* and *Anopheles* mosquitoes, primary vectors of dengue virus and malaria parasites, respectively. *E. anophelis* possesses a broad-spectrum antiviral phenotype, yet a gap in knowledge regarding the mechanistic basis of its interaction with viruses exists.

**Methodology/Principal findings:** To further understand the antiviral interactions between *E. anophelis* and Zika virus (ZIKV), we utilized a non-targeted multi-omics approach, analyzing lipids, proteins, and metabolites of cell monolayers co-infected with ZIKV and *E. anophelis*. We further assessed the gene expression of ZIKV when cultured in the presence of *E. anophelis*. ZIKV cultured in the presence of *E. anophelis* resulted in an attenuated replicative fitness and unproductive virus infection. Further, in this treatment, we observed lower levels of the nonstructural protein 5 (NS5) and RNA-directed RNA polymerase (RdRp) protein. Lastly, a significant decrease in arginine levels, an essential requirement for viral replication and progression of viral infection was observed.

**Conclusions/Significance:** This study provides insights into the molecular basis of *E. anophelis* antiviral phenotype. These findings improve our knowledge of how microbes and viruses interact to impact viral replication. In the future, our findings can be utilized to unravel the mechanism behind the antiviral phenotype of *E. anophelis,* and this can help develop novel paradigms for viral therapeutics.

**AUTHOR SUMMARY:** Zika is a re-emerging disease and is endemic in many regions of sub-Saharan Africa, Asia and Latin America. It remains a major public health threat and lacks FDA-approved therapeutics or vaccines, hence the urgent need for the identification of alternative approaches that limit the transmission of the pathogens by its primary vector, *Aedes* spp. The microbial communities residing in the mosquito midgut play a key role in determining the outcome of mosquito pathogen infection. Flavobacteria dominates the mosquito midgut including *Elizabethkingia*, which is a gram-negative bacillus prevalent in *Aedes* and *Anopheles* species of mosquitoes. *E. anophelis*, a poorly studied midgut microbe, has a broad-spectrum antiviral phenotype, yet the mechanism of its antiviral action is unknown. In this study, we have identified several pathways as well as Zika virus proteins perturbed when the Zika virus is cultivated in the presence of *E. anophelis*. Our findings do not only provide insights into microbial, virus interaction but could be harnessed to develop novel antiviral tools.

## INTRODUCTION

Zika virus (ZIKV) is an emerging virus infection[1–6], endemic in many regions of sub-Saharan Africa [7–15] as well as in countries in South Asia [16–19]. ZIKV is a focus of intense research due to its rapid geographic spread in the Americas and its association with birth defects (e.g. microcephaly) and neurological syndromes[20–22]. It is mainly transmitted by the bite of infected *Aedes* mosquitoes, in addition, sexual, transplacental, as well as transmission through blood transfusion have been documented [23]. There is no FDA-approved medication or vaccine to treat or prevent ZIKV infection despite WHO declaring Zika a public health emergency of international concern [24]. Medications for symptomatic relief are the only source of relief for infected individuals [25]. Control of the mosquito vector is the primary method to limit transmission of ZIKV[23]. However, an upsurge in insecticide resistance as well as changing weather patterns, which can complicate vector control efforts has led to significant geographical range expansion of *Aedes* spp.[26], underscoring the urgent need for novel, efficacious, and cost-effective alternatives to traditional mosquito control, and methods to limit pathogen transmission.

The microbial community residing in the mosquito midgut plays a key role in determining the outcome of the pathogen infection of mosquitoes. The midgut microbiota interferes with pathogen infection[27–34] by activating the basal immunity of the mosquito[29,30,35] and also through their metabolites[36–39]. Due to their potential as candidates for the development of therapeutic and transmission-blocking agents, investigators have studied the interaction of gut microbial taxa and arboviruses of medical importance [28,30,40–46]. *Wolbachia pipientis*, an endosymbiotic bacterium naturally found in 40% of insects has been studied extensively for its anti-viral activity. At the field application level, *Wolbachia* has demonstrated significant success in containing the spread of dengue and Zika in natural populations[47,48]. An expansion of the biological control toolbox by exploration for a wide array of microbes that bear antiviral properties is urgent.

*Elizabethkingia* is a rod-like, gram-negative, aerobic, non-fermenting, non-motile, and non-spore-forming bacteria widely distributed in natural environments, including soil, freshwater, and hospitals[49]. Originally isolated from the midgut of *Anopheles gambiae*[50], it has drawn much attention due to its association with *Anopheles* and *Aedes* spp., primary vectors of *Plasmodium* parasites and dengue virus (DENV), respectively[51]. Due to its initial isolation from the midgut of *Anophelis gambiae*, there is a potential for *E. anophelis* to interact with other co-habiting microbial species in the mosquito midgut and with the host mosquito.

*E. anophelis* was noted as the dominant microbial genus in a lab colony of adult *An. funestus* and *An. arabiensis* collected from Mozambique and South Africa[52]. Further, sequencing of gut bacterial communities of *An. gambiae* from Cameroon in West Africa resulted in 95% of sequence tags associated with *Elizabethkingia spp*. [53]. In addition, *Elizabethkingia* was identified in a Brazilian *Ae. aegypti* colony among the non-fed, blood-fed fed as well as the ZIKV-infected mosquitoes[54]. In our previous findings, we identified *E. anophelis* as the most abundant gut microbiome species in both noninfectious and infectious blood-fed lab-reared *Ae. albopictus* collected from Long Island, New York [55]. A past study reported that 10^7^ *E. anophelis* per midgut, resulted in a significant reduction of *Plasmodium falciparum* oocyst load to approximately zero. Further, the study identified the capacity of *E. anophelis* to inhibit *P. berghei* ookinete development *in vitro*. The study postulated that *E. anophelis* inhibits parasite development either through direct interaction with the parasites or through the production of inhibitory factors[56]. A different study reported that extracts obtained from cultured *E. meningoseptica* showed activity against gram-positive *Staphylococcus aureus* and gram-negative *E. coli* as well as *Candida albicans*. Further, the extracts were active against blood and gametocyte transmission stages of *Plasmodium falciparum*[57]. Consistent with previous studies, we previously showed that *E. anophelis* has a broad-spectrum antiviral activity significantly reducing viral loads of ZIKV, dengue virus (DENV), and chikungunya virus (CHIKV) *in vitro* while negatively impacting infection rates of ZIKV in *Ae. albopictus* mosquitoes[58].

This study aimed to identify the pathways that *E. anophelis* targets to effect, the anti-viral phenotype, as well as delineate the impact of *E. anophelis* on ZIKV gene expression. To address this aim, we cultured ZIKV in the presence of *E. anophelis* and utilized a multi-omics approach, leveraging non-targeted metabolomics, proteomics, and lipidomics assay. The findings of our study will provide insights into the key points targeted by *E. anophelis* to impact ZIKV replication. Further, we contribute to a body of knowledge that can be harnessed to develop new avenues for pathogen and vector control tools.

## METHODS

### Ethics statement

All experiments using viruses or bacteria were performed under biosafety level 2 (BSL-2) conditions at Texas Tech University with Institutional Biosafety Committee approval.

### Cells, Virus, Bacteria

Vero E6 and CCL-81 cell lines were obtained from ATCC. The cell lines were maintained in DMEM (ThermoFisher Scientific, 11965092), supplemented with 10% fetal bovine serum (FBS, Gibco, 16140071) and 1% penicillin/streptomycin (Gibco, 15140148). The cells were incubated at 37°C with 5% CO_2_.

The ZIKV strain utilized in this study (GenBank:KX262887) was obtained through BEI Resources, NIAID, NIH: Zika Virus, R103451, NR-50355. The virus was isolated on January 6, 2016, from the placenta of a human who had traveled to Honduras in 2015. To generate stock for experiments, the virus was passaged three times on Vero E6 cell line to create stock virus and titrated by standard plaque forming assays. Stocks were stored in liquid Nitrogen prior to use.

*Elizabethkingia anophelis*, strain Ag1, NR-50124 (GenBank: CP023402.1) was obtained through BEI Resources, NIAID, NIH. The *E. anophelis* strain Ag1 was isolated in 2010 from the midgut of a mosquito (*Anopheles gambiae*, strain G3) in Las Cruces, New Mexico USA.

### Co-infection of Vero E6 monolayers with bacteria supernatants and Zika virus

Six well plates were seeded with Vero E6 cells three days prior. Upon attaining confluency, the monolayers were inoculated sequentially with 75 μl of 8.0 log_10_ *E. anophelis* CFU/ml and ZIKV at a multiplicity of infection (MOI) of 0.1 (ZIKV/*E. anophelis*); 75 μl of 8.0 log_10_ *E. coli* and ZIKV at an MOI of 0.1(ZIKV/*E. coli*) and mock infected with DMEM maintenance media and ZIKV at an MOI of 0.1(ZIKV/*DMEM*). Infected cells were cultured at 37°C with 5% CO_2_ for 48 hours[55]. Supernatants was harvested from each well, ZIKV RNA was extracted from the supernatants using QIAamp viral RNA kit (Qiagen, 52906) according to the manufacturer protocol.

### Zika virus RNA quantification by RT-qPCR

Quantitative real-time qRT-PCR assay was performed using primers targeting the NS1 region of the ZIKV (ZIKV 1086 CCGCTGCCCAACACAAG; ZIKV 1162C CCACTAACGTTCTTTTGCAGACAT; ZIKV 1107-FAM AGCCTACCTTGACAAGCAGTCAGACACTCAA)[59]. Standard curve was generated using an 8-fold dilution of previously plaque-tittered ZIKV for the quantification of copies of ZIKV genome present in the samples. Plaque assay of a select number of samples were used to confirm the qRT PCR results. The experiment was conducted as six biological and two technical replicates.

### Growth curves

Supernatants harvested from wells inoculated with ZIKV/ *E. anophelis* or ZIKV/ *E. coli* above were filtered using a 0.02-micron filter (Whatman) to exclude the bacteria. Thereafter, the filtered supernatants (ZIKV/*E. anophelis* or ZIKV/*E. coli*) were added to confluent Vero E6 monolayers at equal ratios at 0.1 MOI, ZIKV/*DMEM* was added as a negative control. After incubating at 37°C with 5% CO_2_ for 1 hour, the viral supernatants inoculum was removed and 3 ml of DMEM containing 10% FBS was added. Infected cells were cultured at 37°C with 5% CO_2_ and viral culture supernatants were collected at 24, 48, 72 and 96 hpi time points. Virus titers were determined by RT-qPCR targeting the NS1 region as described above. Standard curves were generated using a 10-fold serial dilution of previously tittered ZIKV for the quantification of copies of the ZIKV genome present in samples.

### Plaque assay

A total of 100 μl of the filtered supernatants of ZIKV/*E. anophelis* and ZIKV/*DMEM* (as control) were serially diluted eight-fold with DMEM containing 1% penicillin/streptomycin. Subsequently, 100 μl of each diluent were added to CCL-81 cell monolayers in six well plates in duplicates and incubated at 37°C with 5% CO_2._ The supernatants containing the virus was discarded after 1 hour of incubation and 3 ml methylcellulose overlay was then added to each well. The infected cells were cultured for another 4 days and then fixed overnight at room temperature by adding 10% formaldehyde solution and stained overnight with 0.5% crystal violet solution. The plaques were counted for the calculation of virus titers.

### RNA-sequencing and data analysis

Total ZIKV RNA was extracted from supernatants harvested after 48 hours from VeroE6 monolayers inoculated with ZIKV/*E. anophelis* or ZIKV/DMEM using RNeasy kit (Qiagen) according to manufacturer’s protocol. The RNA library construction and high-throughput sequencing were performed by Genewiz, Azenta Life Sciences. Multiplexed libraries were sequenced for 150bp at both ends using an Illumina HiSeq6000 platform. Three biological replicates of each condition were RNA sequenced.

The high throughput Illumina sequence data were processed using the Galaxy online tools (usegalaxy.org). Reads were first trimmed and filtered to remove the ambiguous nucleotides and low-quality sequence using trimmomatic (HEADCROP, 15; SLIDING WINDOW, 4, 30; MINLEN, 50). The filtered reads were mapped to the ZIKV reference genome NCBI ViralProj 411812 (RefSeq: GCF_002366285.1), using RNA STAR. ZIKV genome is a polyprotein, and hence, a homogenous distribution of reads was expected along the polyprotein sequence. This was not the case in this study and hence the annotation of the unique coding sequence of the genome was split into the unique proteins coded within the polypeptide. The new annotation generated was used to count reads aligning uniquely to the specific genes with htseq count. Subsequently, differential expression (DE) was calculated using the reads aligning uniquely to ZIKV transcripts using DESeq2 [60]. Significance was determined using DESeq2 exact test for the negative binomial distribution, corrected with a False Discovery Rate (FDR) at P < 0.05.

### Untargeted analysis of proteins, lipids and metabolites

Confluent monolayers of Vero E6 were infected with ZIKV and DMEM (ZIKV/DMEM); ZIKV and heat inactivated *E. anophelis* (ZIKV/Heat inactivated *E. anophelis* ) or ZIKV and live E. anophelis (ZIKV/live *E. anophelis*). The samples were cultured at 37°C with 5% CO_2_ for 48 hours. At 48 hours post infection (hpi), supernatants were harvested from wells containing ZIKV/DMEM, ZIKV/Heat inactivated and ZIKV/live *E. anophelis.* To assess whether ZIKV replication was successful, ZIKV RNA was extracted from the supernatants using QIAamp viral RNA kit (Qiagen, 52906) according to manufacturer protocol. Samples were stored at -80°C before submission to the Center for Biotechnology and Genomics at Texas Tech University for untargeted screening of proteins, lipids and metabolites. Microfuge tubes handled by the individual involved in the sample preparations were sent to the sequencing core for normalization of the variation introduced during sample collection and preparation. The maintenance media that was utilized to maintain the cells was collected as blank negative control. The samples were pooled into four pools with six biological replicates [ZIKV/DMEM (Mock infection); ZIKV/Heat inactivated *E. anophelis* and ZIKV/live *E. anophelis*].

To characterize the protein changes that occur when ZIKV is co-cultured with *E. anophelis*, we performed protein analysis on the four pools by BCA protein assay. Based on the protein concentrations, aliquots of the four samples containing 50 μg proteins were aliquoted into a separate tube and volume was adjusted to 50μl using 50mM ammonium bicarbonate buffer solution. Proteins were denatured at 90°C for 15 min and subsequently reduced by adding 200mM of DL-Dithiothreitol (DTT) by incubation at 60°C for 45 min. Following protein reduction, 200mM of Iodoacetamide (IAA) and incubated at 37°C for 45 min to allow protein alkylation. DTT was added a second time to quench the extra IAA and incubated at 37°C for 30 mins. An aliquot of 2 μg of trypsin was added to each sample (trypsin: protein=1:25, m/m) and incubated at 37°C for 18 h. After tryptic digestion, formic acid (FA) was added to the final concentration of 1% (v/v) to stop the digestion process. Samples were dried in a speed vacuum, then resuspended to 1 μg/μL with a solution of 20% acetonitrile (ACN) and 1% FA. After resuspension, 1 μl of each sample (1 μg of proteins) was injected and analyzed by LC-MS/MS (Thermo UltiMate 3000 nanoLC/ Thermo Fusion Lumos). The protein identification and quantification were performed by Thermo Proteome Discoverer 2.4 by querying the database of Chlorocebus Sabaeus (Green Monkey) acquired from Uniprot https://www.uniprot.org/proteomes/UP000029965, the proteins of ZIKV (GenBank:KX262887) and *Elizabethkingia anophelis*, strain Ag1, NR-50124 (GenBank: CP023402.1). The statistical significance (*p*<0.05) of protein expression changes was identified after performing the Mann-Whitney *U* test.

To analyze the Metabolites and Lipids, a volume of 200 μl of extraction solution [dichloromethane (DCM): methanol=1:2 (v/v)] was added to 100μl of the sample aliquots above, vortexed for 30 seconds and incubated on ice for 45 min. A total volume of 75 μl of DCM was added to the solution after incubation and vortexed for 30 s followed by addition of 75 μl of cold water, vortex for 30 s and centrifuged at 5000 rpm for 15 min. The top layer (aqueous phase), which contained polar/hydrophilic metabolites, was collected for metabolomics analysis. The bottom layer (organic phase), which contained non-polar/hydrophobic lipids, was collected for lipidomics analysis. Both the aqueous layer and organic layers were air dried, then resuspended before LC-MS/MS analysis.

Metabolomics samples were resuspended to a volume of 50μl using a 1:1 solution of 50% methanol, 50% water while Lipidomics samples were resuspended in a volume of 50μl of a solution of 65% ACN, 30% Isopropanol, 5% water. Samples were analyzed using Thermo Vanquish UHPLC/ Thermo Q Exactive HF. After running pool samples for tests, the injection volume for metabolomics samples was decided as 5μl, whereas lipidomics samples as 10μl. The identification and quantification of metabolites and lipids were performed by Thermo Compound Discoverer 3.1 and LipidSearch 4.2, respectively. The metabolites/lipids with significant changes (*p*<0.05) were identified after performing the Mann-Whitney *U* test.

### Phenylalanine and Arginine assay

We observed a reduction in levels of phenylalanine and arginine among the ZIKV/*E. anophelis* samples in the metabolomic assay. To confirm changes in phenylalanine and arginine concentration upon culturing ZIKV in the presence of *E. anophelis*, a secondary biochemical assay was performed to verify the metabolomic predictions. Both analyses were performed using commercial assay kits.

Phenylalanine assay (Sigma-Aldrich MAK005) utilizes a coupled enzyme assay that results in the deamination of phenylalanine and the production of NADH which reacts with the probe resulting in a fluorescent (λ_ex_ = 535 nm/λ_em_ = 587 nm) product, proportional to the phenylalanine present.

Confluent VeroE6 monolayers were inoculated with ZIKV/DMEM or ZIKV/*E. anophelis* and cultured for 48 hours. A total of 50μl supernatant was harvested at 48 hpi and filtered in a 10kDa MWCO spin filter (Sigma-Aldrich) by centrifuging at 13 000 g for 10 minutes to remove insoluble material. Samples were diluted 1:100, phenylalanine assay buffer, enzyme mix, and developer were mixed together, standards and sample blank were run in duplicates, this was mixed, incubated for 20 minutes at 37°C. The fluorescence intensity was measured at (λ_ex_ = 535 nm/λ_em_ = 587 nm. A standard curve was generated and the amount of phenylalanine present in the ZIKV/DMEM and ZIKV/*E. anophelis* supernatant was determined from the standard curve.

L-Arginine assay (Sigma-Aldrich MAK370) is an enzyme-based assay whereby L-Arginine is converted into a series of intermediates which reacts with the probe resulting in a stable colorimetric signal at 450 nm (A_450_). A total of 2 μl of sample cleanup mix was added to 100 μl of ZIKV/DMEM and ZIKV/*E. anophelis* supernatants, incubated at 37°C for 1 hour and centrifuged in Corning Spin-X UF concentrator (Corning) at 13 000 g for 10 minutes at 4°C. Arginine enzyme mix, assay buffer was added to the samples and standards and incubated at 37°C for 30 minutes. The reaction mix containing probes was then added and incubated at 37°C for 60 minutes. The assay was run in duplicates. The absorbance was measured at 450nm. The amount of arginine present in the ZIKV/DMEM and ZIKV/*E. anophelis* supernatant was determined from the standard curve.

### Statistical analysis

Data analysis was carried out using GraphPad Prism software. Statistical evaluation was performed using one-way ANOVA or student’s unpaired t-test for single factor analysis. Unless stated otherwise the experiments were done in six biological replicates and two technical replicates. The data are presented as means.

## RESULTS

### *Elizabethkingia anophelis* attenuates Zika virus replicative fitness and yield of infective virus

We have previously demonstrated the antiviral effect of *E. anophelis* on ZIKV, DENV, and CHIKV[58]. In the current study, ZIKV cultured in the presence of *E. anophelis* resulted in a significant reduction of ZIKV RNA copy numbers (unpaired t-test; P< 0.0001) (Figure 1A) but this was not the case when cultured in the presence of *E. coli* (unpaired t-test; P = 0.4947) (Figure 1B);(Supplementary file no. 1).

**Figure 1.**
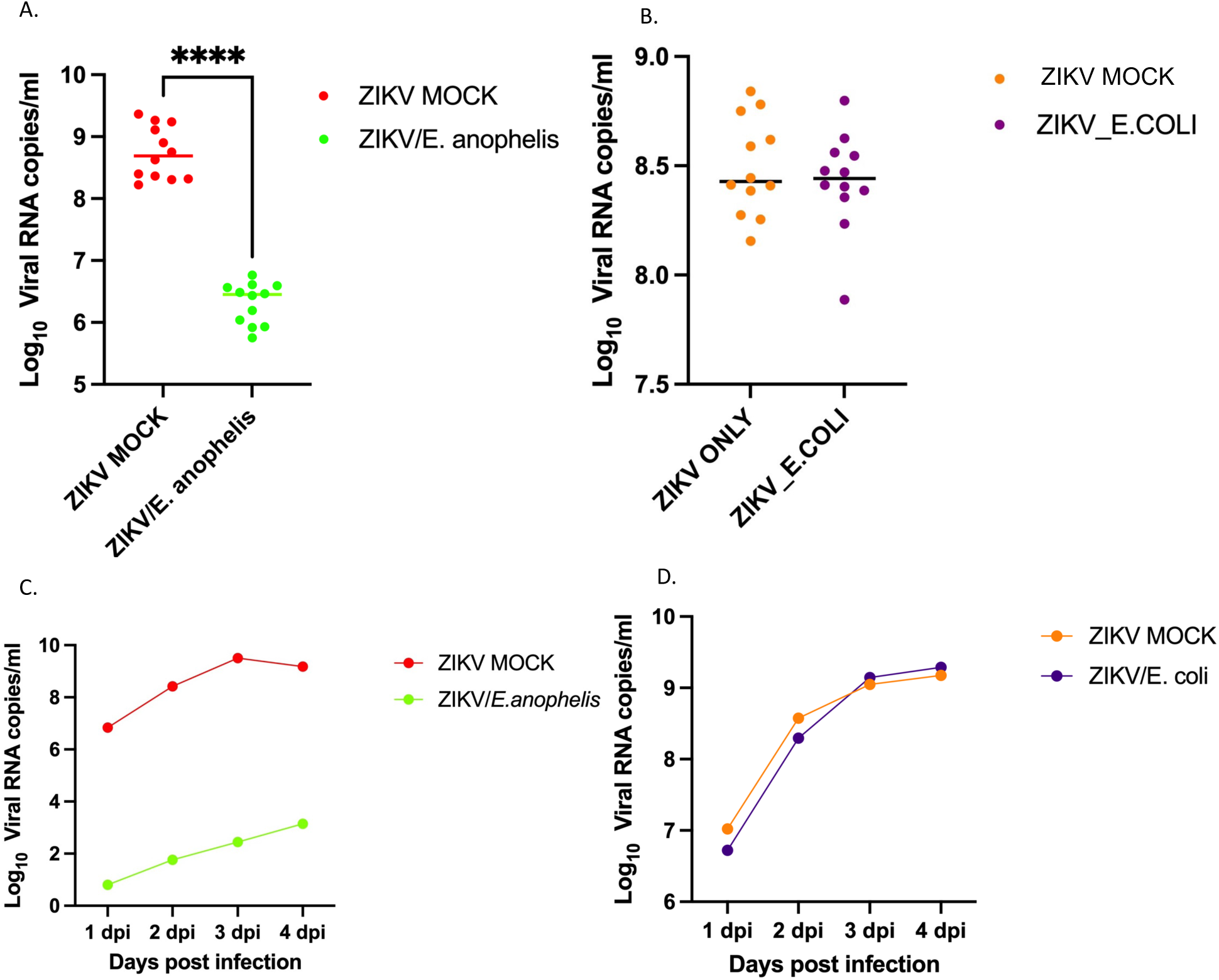
Analysis of the impact of *E. anophelis* on viral replication and growth kinetics of ZIKV. Vero E6 monolayers were sequentially inoculated with either Zika virus (ZIKV) and live *E. anophelis* or Zika virus and *E. coli.* Monolayers mock infected with DMEM and ZIKV were the negative controls. At 2 dpi, the viral supernatant was harvested, and the virus was quantified by qRT-PCR. **A.** The viral supernatants harvested from cell monolayers were sequentially inoculated with ZIKV*/E. anophelis* demonstrated an attenuated replication (unpaired t-test; P< 0.0001) compared to **B.** ZIKV/*E. coli* (unpaired t-test; P = 0.4947). ZIKV*/E. anophelis* or ZIKV/*E. coli* supernatants harvested at 48 hpi were filtered through a 0.2µm filter to preclude bacteria, ZIKV/DMEM was the negative control. A growth curve assay was performed utilizing RT-qPCR targeting the ZIKV NS1 region to quantify the ZIKV RNA copy numbers at 1, 2, 3, and 4 dpi. **C**. ZIKV/*E. anophelis* samples demonstrated an attenuated replicative fitness at every time point while **D.** ZIKV/*E. coli* samples did not show any alteration in its growth patterns and the replication was similar at every time point. The virus genome copy number values are means standard deviations (representative experiment of three biological replicates of two independent experiments).

Next, we queried whether culturing ZIKV in the presence of *E. anophelis* would impact virus replicative fitness. To preclude the effect of the bacteria, we filtered out the bacteria (*E. anophelis* or *E. coli*) from the viral supernatants before performing the growth curve assay. ZIKV exposure to *E. anophelis* resulted in subsequent suppressed replication throughout the growth curve assay (Figure 1C; Supplementary file 2). We observed a significant variation in the ZIKV RNA copy numbers between ZIKV/*E. anophelis* and ZIKV/DMEM (mock infection) samples at each time point. The ZIKV/DMEM (mock infection) demonstrated increased replication relative to ZIKV/*E. anophelis* at all time points and, at 5 dpi, there was an approximately a two-fold difference in RNA copy numbers between the two treatments. In contrast, we did not measure any difference in genome copy numbers in ZIKV/DMEM and pre-filtered ZIKV/*E. coli* samples at each time point throughout the growth curve assay (Figure 1D; Supplementary file 2).

Next, we assessed the impact of *E. anophelis* on the formation of infectious virus. As above, we cultured ZIKV in the presence of *E. anophelis*, and harvested ZIKV/DMEM and ZIKV/*E. anophelis* supernatants at 48 hpi, filtered the bacteria, and seeded confluent CCL-81 with serial dilutions up to 10^-8^. We did not observe any plaque formation in any well in replicate plates inoculated with filtered ZIKV/*E. anophelis* (Figure 2). The control wells inoculated with ZIKV/DMEM supernatants had visible plaques up to 4^th^ dilution in one plate and the 5^th^ dilution in the replicate plate (Figure 2).

**Figure 2.**
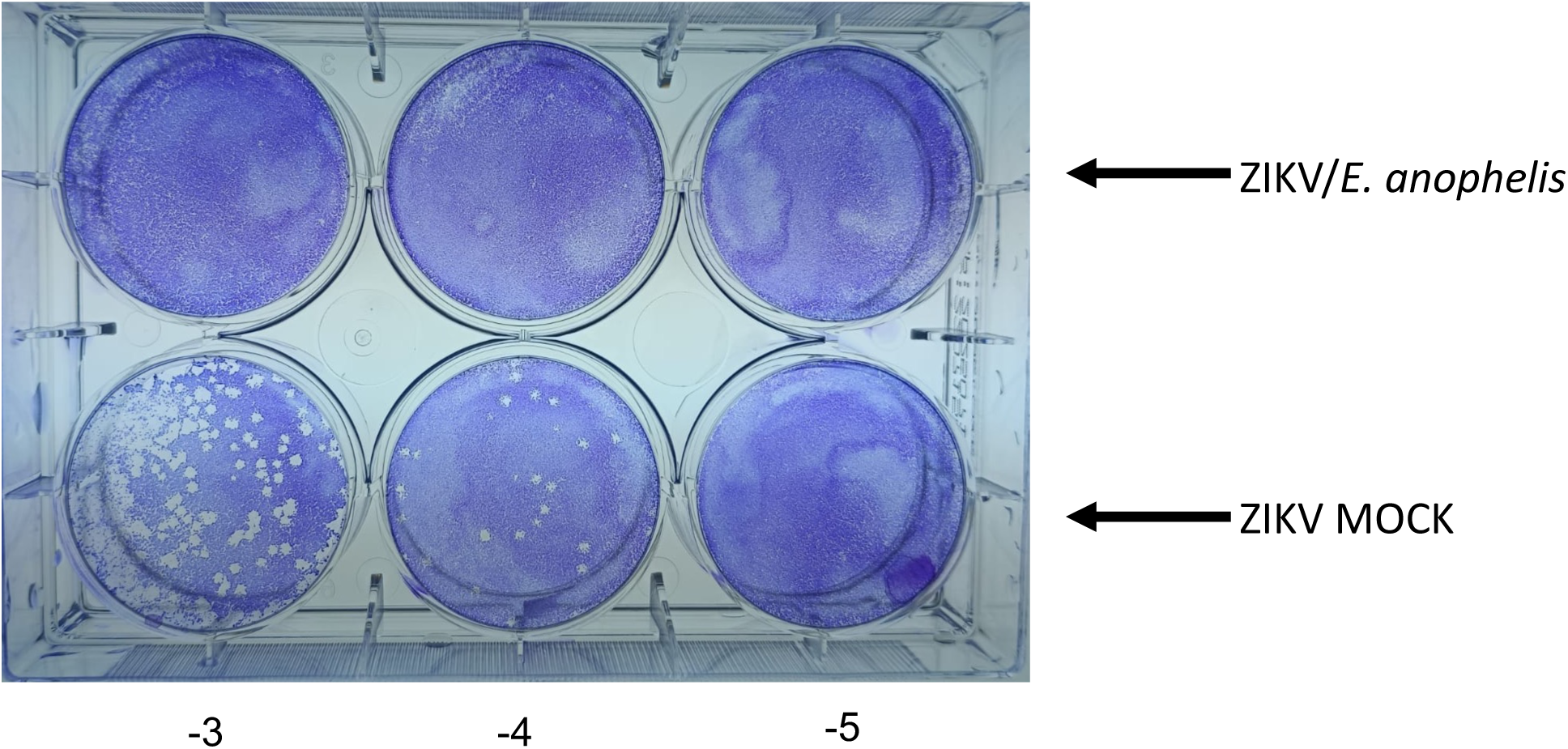
Determination of the impact of *E. anophelis* on ZIKV infectiousness. ZIKV*/E. anophelis* supernatants harvested at 48 hpi were filtered through a 0.2µm filter to preclude bacteria, and ZIKV/DMEM was used as a negative control. A total of 100μl of each supernatant was used to inoculate Vero CCL-81 cell monolayers. ZIKV*/E. anophelis* demonstrated a loss in capacity to form an infective virus.

### Reduction of Zika virus NS5 levels

We utilized transcriptomic analysis to investigate ZIKV gene expression responses to *E. anophelis*. We carried out an RNA-seq analysis of supernatants originating from Vero E6 monolayers inoculated with ZIKV/*E. anophelis* and ZIKV/DMEM (mock infection). We observed a statistically significant reduction of the ZIKV nonstructural protein 5 (NS5) levels in ZIKV/*E. anophelis* supernatants (Supplementary file 3) (P=0.0004). Further, both the membrane and pre-membrane genes showed a minimal increase in levels. Additionally, though not statistically significant, we observed a decrease in the amount of the envelope, NS1, NS2B, NS3, and NS4A genes. In addition, when we queried the proteins identified in the multi-omics analysis against ZIKV polyprotein we observed a statistically significant increase in the relative abundance of pre-membrane (prM), envelope protein (E), and nonstructural protein 3 (NS3) among the ZIKV/*E. anophelis* and ZIKV/Heat inactivated *E. anophelis* samples (Supplementary Figure 1).

### Global protein, lipid, and metabolite marker isolation

A total of 21 samples, 18 supernatant samples [ZIKV/live *E. anophelis*, n = 6; ZIKV/Heat inactivated *E. anophelis*, n = 6 and ZIKV/DMEM, n=6] with an average viral RNA copy number range of Log_10_ 7.4 – 8.2 RNA copies/ml and a standard deviation range of ±0.13 – 0.31(Supplementary file 4) and three negative controls made up of maintenance media only (DMEM and 10% FBS) were subjected to untargeted proteomic, lipidomic and metabolomic analysis. Globally, 403 lipids(Supplementary file 5), 115 proteins (Supplementary file 6), and 145 metabolites (Supplementary file 7) with statistically significant changes (*p*<0.05) were identified.

### Significant reduction of proteins responsible for replication of RNA-based viruses

Analysis of the unsupervised principal component analysis (PCA) revealed a unique cluster of proteins associated with ZIKV/live *E. anophelis* samples, a few proteins shared among ZIKV/live *E. anophelis* and ZIKV/heat-inactivated *E. anophelis* conditions, while ZIKV/DMEM and ZIKV/Heat inactivated *E. anophelis* samples shared common proteins (Figure 3A).

**Figure 3.**
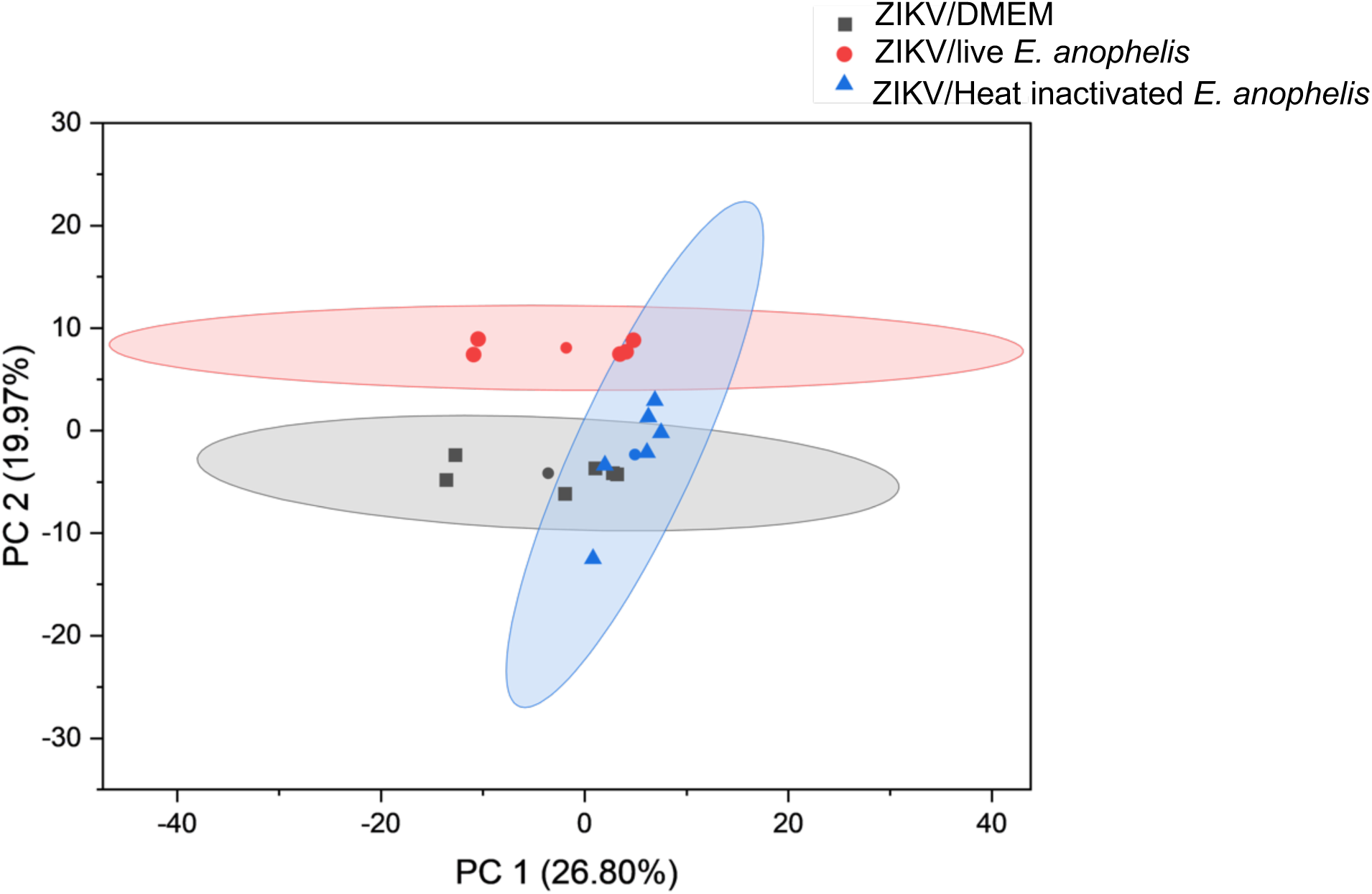
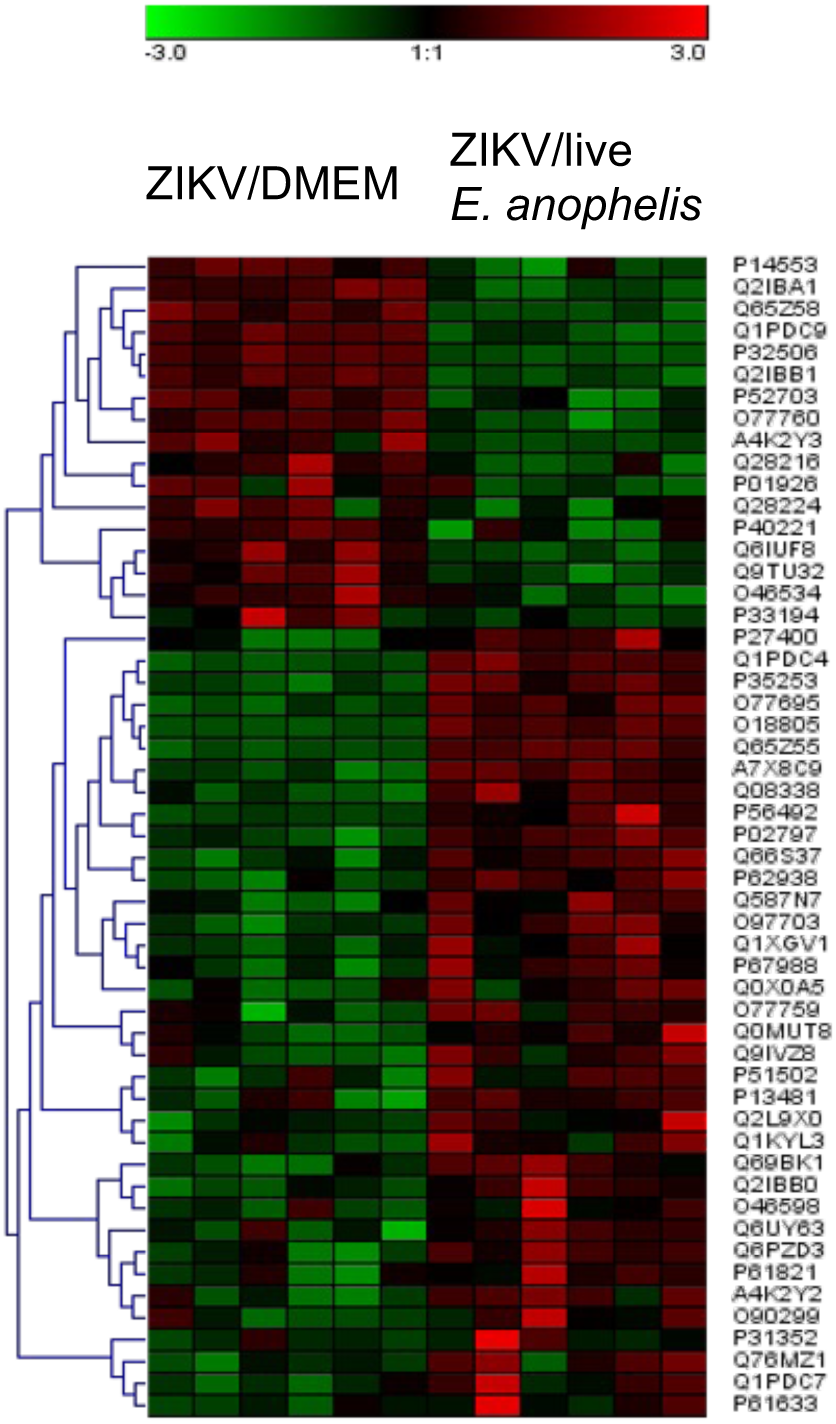
Assessment of *E. anophelis* on protein profile. **A**. Analysis of the unsupervised principal component analysis (PCA) revealed a unique cluster of proteins associated with ZIKV/live *E. anophelis* samples, a few common proteins between ZIKV/live *E. anophelis* and ZIKV/Heat inactivated *E. anophelis* samples, while ZIKV/DMEM and ZIKV/Heat inactivated *E. anophelis* samples shared common proteins. **B**. An assortment of proteins associated with viral replication was impacted in the ZIKV/live *E. anophelis* samples. Point coloration represents the fold change observed in ZIKV/*E. anophelis* samples.

We observed a significant reduction in the levels of RNA-directed RNA polymerase (RdRp) protein among the ZIKV/live *E. anophelis* samples. RdRp is important in the replication of RNA-based viruses[61] (Figure 3B). In addition, a reduction in polymerase cofactor VP35, a viral structural protein which apart from inhibiting host innate immune response, acts as a cofactor in RNA polymerase transcription and replication complex cofactor was evident among the ZIKV/live *E. anophelis* samples. Gag polyprotein known to drive the formation of virions from productively infected cells was significantly reduced among the ZIKV/live *E. anophelis* samples.

### Perturbations of lipids associated with membrane permeability

In this study, ZIKV/live *E. anophelis* had a distinct lipid profile uncommon to ZIKV/DMEM and ZIKV/Heat inactivated *E. anophelis* which shared common lipids (Figure 4A).

**Figure 4.**
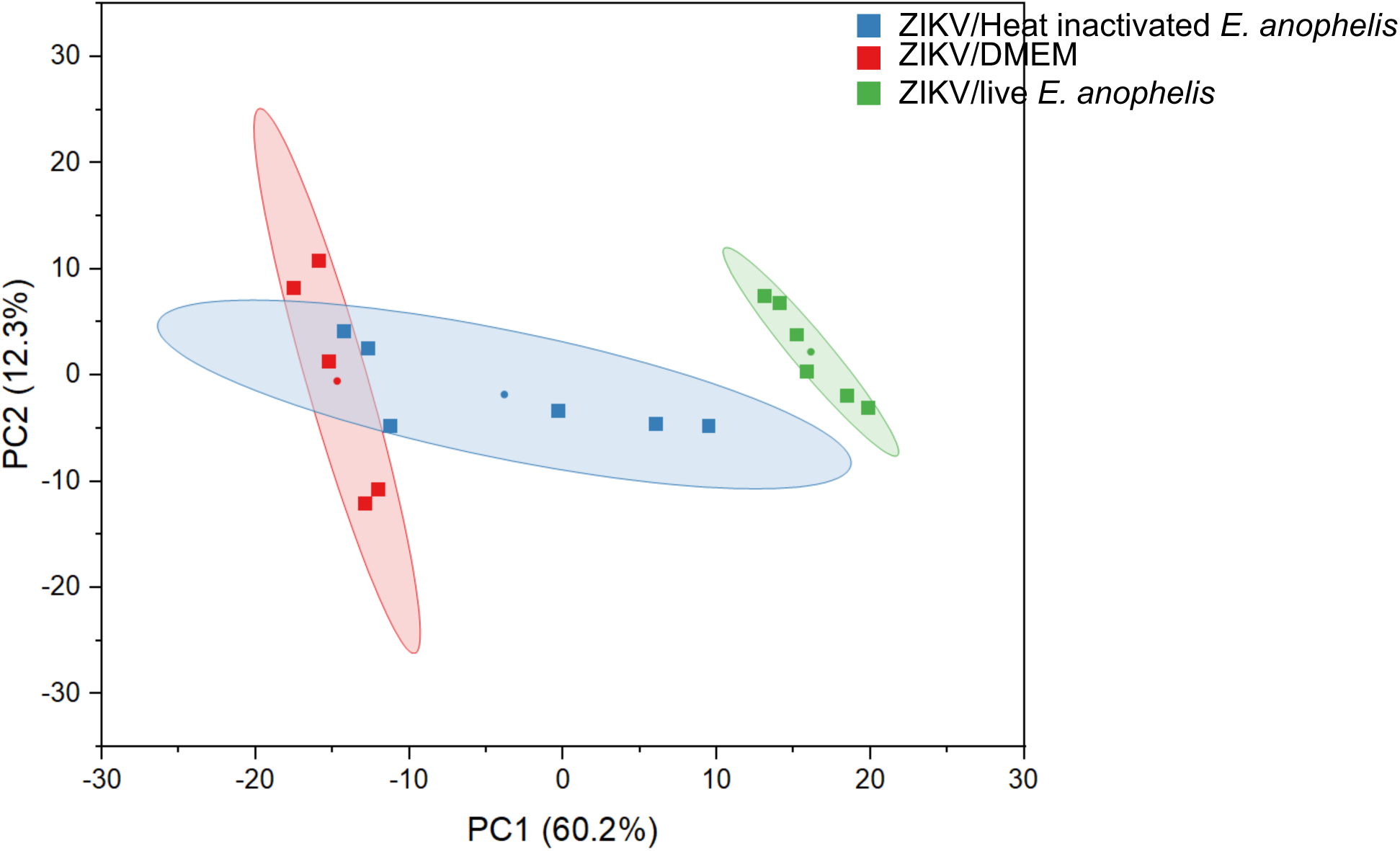
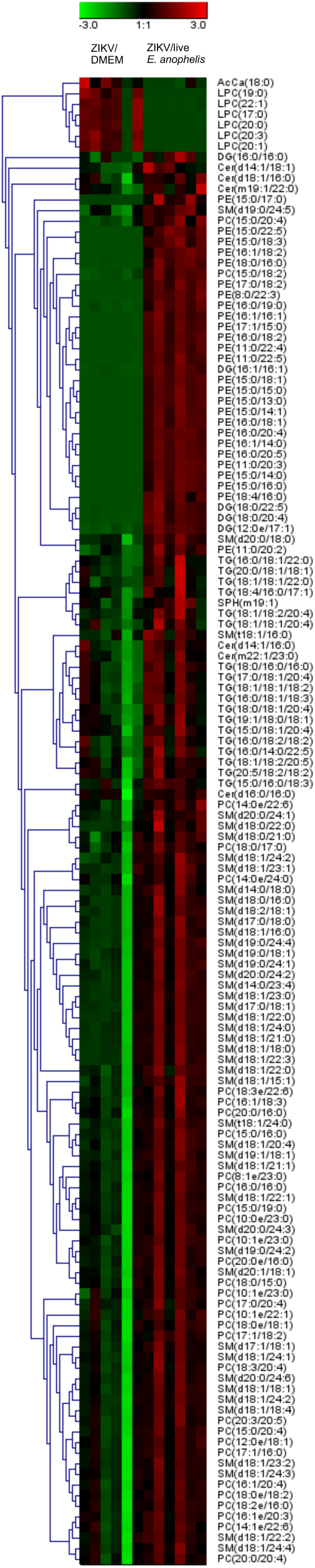
Defining the repertoire of host lipids perturbed by *E. anophelis*. **A.** The lipid profile demonstrated distinct lipids associated with ZIKV/ live *E. anophelis* and that differed from ZIKV/DMEM and ZIKV/Heat inactivated *E. anophelis* **B.** Interaction of ZIKV and *E. anophelis* results in deviations associated with lipids associated with membrane curvature such as Lysophosphatidylcholine and fatty acids associated with formation of viral replication factories. We also observed the upregulation of lipids such as sphingomyelin and ceramide, that are known to be associated with the viral membrane entry. Point coloration represents the fold change observed in ZIKV/*E. anophelis* samples.

Analysis of the lipid profile demonstrated multiple deviations, in particular, lysophosphatidylcholine, a lysolipid, known to promote positive curvature and enhance membrane permeability was reduced among ZIKV/live *E. anophelis* samples. All ZIKV/live *E. anophelis* samples except one, showed a reduction of Acyl Carnitine, an ester of I-carnitine and fatty acids. The rest of the lipid profiles identified in this study increased levels: diglycerides, ceramides, phosphatidylethanolamines, sphingomyelins, phosphatidylcholines, and triglycerides (Figure 4B).

### Reduction of metabolites necessary for viral replication

Among the ZIKV/live *E. anophelis* samples, we observed a reduction of arginine and phenylalanine levels known to be important in several viral replication processes. We further validated the changes in phenylalanine and arginine concentration through a secondary biochemical assay which demonstrated a reduction of both arginine (Supplementary file 8) and phenylalanine levels(Supplementary file 9) in supernatants associated with ZIKV/live *E. anophelis* relative to ZIKV/DMEM (mock infection).

Furthermore, among the ZIKV/live *E. anophelis,* there was a decrease of arachidonic acid, a precursor of eicosapentaenoic acid, present in the cell membranes of most body cell samples. Additionally, ZIKV cultured in the presence of *E. anophelis* resulted in a hypoglycemic condition with low sucrose levels observed. Reduced levels of cytidine were detected among ZIKV/live *E. anophelis* samples. Cytidine is a component of RNA and a nucleoside formed when cytosine is attached to a ribose ring. Lastly, Pyridoxal, the natural form of vitamin B6, a co-enzyme involved in the metabolism of carbohydrates, lipids, proteins, and nucleic acids was reduced in the ZIKV/live *E. anophelis* samples (Figure 5).

**Figure 5.**
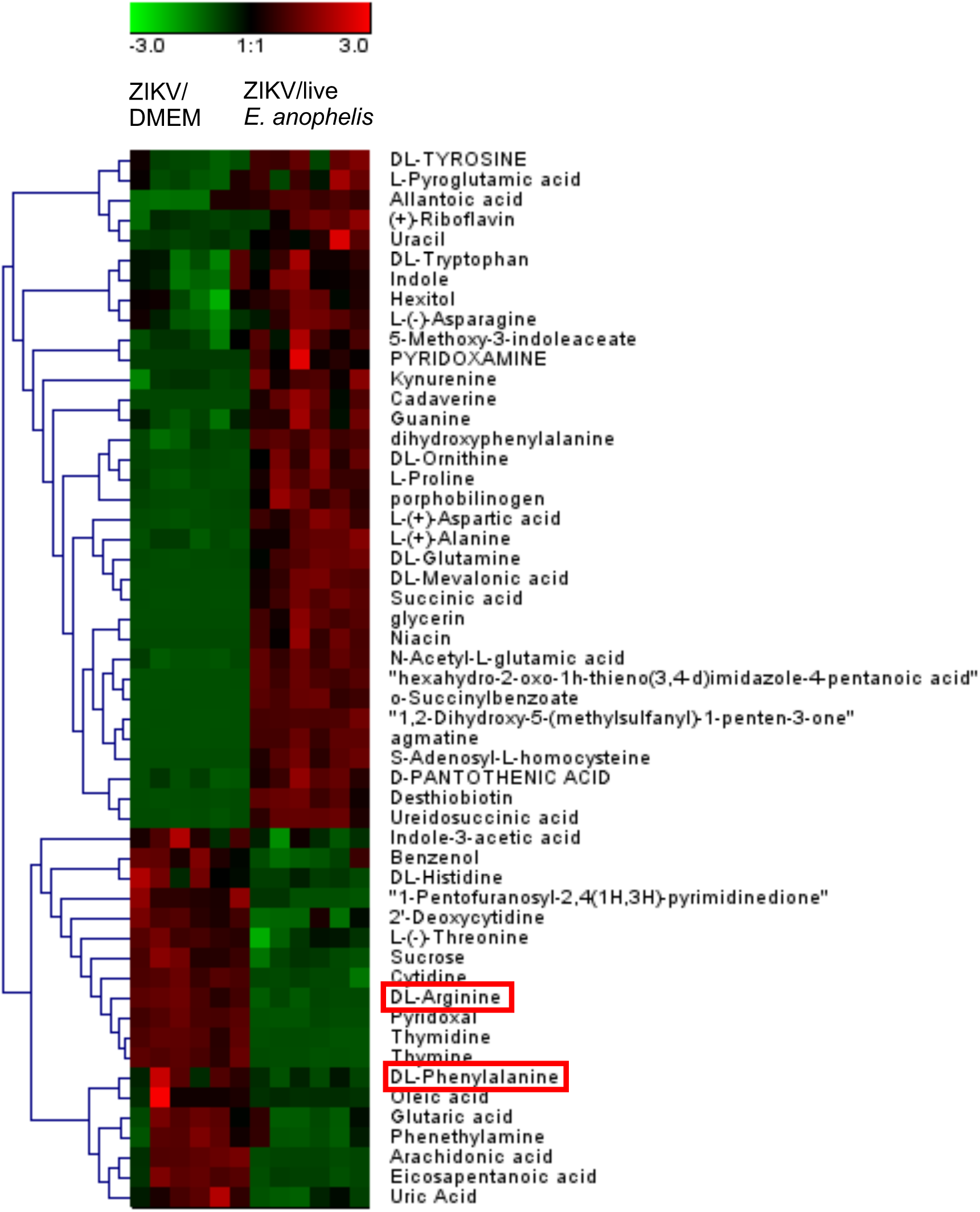
Assessment of the impact of *E. anophelis* on metabolites. We measured a significant decrease in levels of Arginine in ZIKV/ live *E. anophelis* compared to ZIKV/DMEM (Mock). Arginine is an essential requirement for the replication of viruses and the progression of viral infection. Point coloration represents the fold change observed in ZIKV/*E. anophelis* samples.

## DISCUSSION

The results of our study demonstrate that ZIKV cultured in the presence of *E. anophelis* is associated with attenuated viral replicative fitness and lack of infectious virus. Indeed, the lack of robust viral replication in the ZIKV/ live *E. anophelis* samples is coupled with a reduction in levels of NS5 expression and low levels of RdRp protein. The NS5 protein is essential for the replication of the flaviviral RNA genome[62–64]. NS5 nonstructural protein consists of the N-terminal region made up of methyltransferase (MT), this is followed by a short linker that joins the MT to the RdRp[65–67]. Apart from adding the 5’ RNA cap structure to assist in the translation of the viral polyprotein and counteracting antiviral signaling [68,69], the RdRp initiates RNA synthesis [70]. The alteration of NS5 has been shown to affect the speed and fidelity of RNA replication, hence altering the viral fitness[71–74]. We cannot rule out that the variation of NS5 levels observed in our study could also reflect low read counts, technical bias, and ribosome stalling at specific sites [75]. Further investigations involving experimental validation are required as well as cultures on broad cell lines to assess the potential of host-specific antiviral responses.

Flavivirus membrane proteins play important roles in facilitating the entry of viruses into host cells and assembly [76]. After the flavivirus maturation, both prM/E proteins undergo conformational changes resulting from a low pH environment, leading to cleavage of prM into pr and M resulting in a mature virus that can proceed to exit the cell through the secretory pathway[77–79]. In this study, we observed increased levels of prM and E proteins in the ZIKV/ live *E. anophelis* samples relative to ZIKV/DMEM (mock infection). This may suggest the increased utility of prM and E proteins in generating mature virions in ZIKV/DMEM compared to the ZIKV/ live *E. anophelis* samples. Future studies aimed at dissecting what point of the ZIKV life cycle is impacted by *E. anophelis* are important to understand the basis of *E. anophelis* antiviral phenotype.

The significant reduction of both arginine and phenylalanine levels in ZIKV/live *E. anophelis* samples relative to ZIKV/DMEM (mock infection) is interesting. Arginine is an essential requirement for the replication of viruses and the progression of viral infection[80]. The result of our study corroborates previous findings which demonstrated the impact of nutritional deficiencies on Herpes Simplex Virus (HSV) replication in human cells by eliminating arginine, resulting in no viral recovery from any tube containing arginine-deficient medium, with no cytopathic effect (CPE). When the arginine-deficient medium was replaced with a complete medium the CPE was recovered, and viral replication was restored[81]. Further, arginine has been demonstrated to be essential for the replication of vaccinia virus in HeLa cells and the yield of infective virus was equivalent to the concentration of arginine in the medium. It was further shown to be important for RNA and DNA replication and was incorporated and utilized in the formation of complete virus particles. The study concluded that the requirement of arginine in the vaccinia replication cycle can be distinguished as required before the synthesis of virus DNA and associated with the formation of mature virus particles[80].

In this study, all ZIKV/live *E. anophelis* samples except one, showed reduced levels of Acyl Carnitine, an ester of I-carnitine and fatty acids. A previous study focused on identifying the specific cellular pathways and genes necessary for DENV replication demonstrated that fatty acid synthesis is critical for DENV replication and upon inhibition of fatty acid synthesis pathways, a significant inhibition of DENV replication was observed [82]. Further, in the same study, the inhibition of the fatty acid biosynthesis inhibited the replication of the Yellow Fever virus and West Nile virus[82].

Lipid profiles identified in this study were enriched: diglycerides, ceramides, phosphatidylethanolamines, sphingomyelins, phosphatidylcholines, and triglycerides. Our results coincide with the findings of a previous study aimed at profiling the lipidome of DENV-infected mosquito cells at various infection stages. At the binding and entry stage, the study utilized an inactivated virus that was capable of binding and entry but not replication, and they observed selective enrichment of lipids that can influence membrane structure and also have signaling functions such as sphingomyelin, ceramide, lysophospholipids and several intermediates such as mono- and diacylglycerol and phosphatic acid[83]. Taken together, our findings suggest that the lipid profile perturbations in this study may point to virus interaction with the cellular membranes, but a lack of a robust viral replication as demonstrated by a downregulation of cellular components associated with the formation of viral replication factories.

In conclusion, this study has provided insights into the pathways as well as viral genes that are impacted by ZIKV and *E. anophelis* interaction. These results contribute to understanding the molecular determinants of *E. anophelis* antiviral phenotype.

One of the significant limitations of the study is the lack of inclusion of an *in vivo* analysis of the interaction between ZIKV and *E. anophelis* or a comparison of the findings of Vero cell lines with mosquito or human cell lines which could be different and alter our findings. In the future, to build on our study findings, we recommend an *in vivo* study and a broad comparison of different cell lines to gain a basic understanding of whether *E. anophelis* antiviral effects are host specific.

Despite these limitations, our findings provide evidence that a poorly studied midgut microbe, *E. anophelis*, has an antiviral phenotype, reducing the replicative fitness of ZIKV and formation of infectious virus, and is associated with perturbance of pathways associated with key pathways that are necessary for viral replication and formation of viral factories. Future studies to identify the mechanisms utilized by *E. anophelis* to negatively impact viral replication can build upon our current findings. Identification of such mechanisms can lead to novel anti-viral pathways.

## ACKNOWLEDGEMENTS

This publication was supported by start-up funds from Texas Tech University. Its content is solely the responsibility of the authors and doesn’t necessarily represent the official views of Texas Tech University.

## AUTHOR CONTRIBUTIONS

OS: Conducted the experiments and participated in the writing of the manuscript.

WJ: Conducted the experiments, was involved in data analysis, and participated in the writing of the manuscript.

SM: Conducted the experiments and participated in the writing of the manuscript.

RJ: Conducted the experiments and participated in the writing of the manuscript.

ONR: Conducted the experiments and participated in the writing of the manuscript.

GM: Involved in data analysis and participated in the writing of the manuscript.

JP: Involved in data analysis and participated in the writing of the manuscript.

WP: Involved in data analysis and participated in the writing of the manuscript.

NJ: Involved in data analysis and participated in the writing of the manuscript.

CLB: Participated in the writing of the manuscript.

VA: Involved in data analysis and writing of the manuscript.

CAT: Involved in the study design and participated in the writing of the manuscript.

KLD: Involved in the study design and participated in the writing of the manuscript.

MY: Involved in the study design and participated in the writing of the manuscript.

MGO: Involved in the study design, analyzed data, conducted the experiments, and drafted the manuscript.

## Supplementary Materials

### Supplementary figure

**Supplementary figure 1.** Bar plots showing protein expression of ZIKV polyprotein.

### Supplementary files

**Supplementary file 1**. An excel file showing the RT-qPCR results of *E. anophelis* antiviral activity.

**Supplementary file 2**. An excel file showing the RT-qPCR results of ZIKV/*E. anophelis* and ZIKV/*E. coli* growth curve assay.

**Supplementary file 3**. Transcriptomics results demonstrating the impact of *E. anophelis* on ZIKV gene expression.

**Supplementary file 4**. An excel file showing the RT-qPCR results of ZIKV/*E. anophelis* and ZIKV/*E. coli* samples utilized in the multi-omics analysis.

**Supplementary file 5**. An excel file showing the list of lipids identified in this study.

**Supplementary file 6**. An excel file showing the list of proteins identified in this study.

**Supplementary file 7**. An excel file showing the list of metabolites identified in this study.

**Supplementary file 8**. An excel sheet showing the arginine assay results.

**Supplementary file 9**. An excel sheet showing the phenylalanine assay results.

## Notes

### Competing Interest Statement

The authors have declared no competing interest.

